# Crystal structure of the *Arabidopsis* SPIRAL2 C-terminal domain reveals a p80-Katanin-like domain

**DOI:** 10.1101/2022.12.28.522140

**Authors:** Derek L. Bolhuis, Ram Dixit, Kevin C. Slep

## Abstract

Epidermal cells of dark-grown plant seedlings reorient their cortical microtubule arrays in response to blue light from a net lateral orientation to a net longitudinal orientation with respect to the long axis of cells. The molecular mechanism underlying this microtubule array reorientation involves katanin, a microtubule severing enzyme, and a plant-specific microtubule associated protein called SPIRAL2. Katanin preferentially severs longitudinal microtubules, generating seeds that amplify the longitudinal array. Upon severing, SPIRAL2 binds nascent microtubule minus ends and limits their dynamics, thereby stabilizing the longitudinal array while the lateral array undergoes net depolymerization. To date, no experimental structural information is available for SPIRAL2 to help inform its mechanism. To gain insight into SPIRAL2 structure and function, we determined a 1.8 Å resolution crystal structure of the *Arabidopsis thaliana* SPIRAL2 C-terminal domain. The domain is composed of seven core α-helices, arranged in an α-solenoid. Amino-acid sequence conservation maps primarily to one face of the domain involving helices α1, α3, α5, and an extended loop, the α6-α7 loop. The domain fold is similar to, yet structurally distinct from the C-terminal domain of Ge-1 (an mRNA decapping complex factor involved in P-body localization) and, surprisingly, the C-terminal domain of the katanin p80 regulatory subunit. The katanin p80 C-terminal domain heterodimerizes with the MIT domain of the katanin p60 catalytic subunit, and in metazoans, binds the microtubule minus-end factors CAMSAP3 and ASPM. Structural analysis predicts that SPIRAL2 does not engage katanin p60 in a mode homologous to katanin p80. The SPIRAL2 structure highlights an interesting evolutionary convergence of domain architecture and microtubule minus-end localization between SPIRAL2 and katanin complexes, and establishes a foundation upon which structure-function analysis can be conducted to elucidate the role of this domain in the regulation of plant microtubule arrays.

## Introduction

Microtubules are polarized cytoskeletal polymers of the αβ-tubulin heterodimer that undergo dynamic instability (1,2). Microtubules are critical for cellular support and the asymmetric localization of cellular factors either through polarized microtubule motor-dependent transport, or via factors that specifically bind the microtubule plus or minus end. Collectively, asymmetric functions are best achieved when microtubules are arranged in an array that can adapt and reorient in response to intrinsic (e.g. cell cycle regulators) or extrinsic (e.g. a chemoattractant) cues. While some organisms use centrosomes to organize microtubule arrays, many organisms and cell types form acentrosomal microtubule arrays. How these arrays form, are maintained over time, and morph or reorient in response to cues is poorly understood. Higher plants form acentrosomal cortical interphase microtubule arrays that aid in the asymmetric localization of cell wall biosynthesis machinery, a process critical for anisotropic growth and development (3–6). In many tissues, plant acentrosomal microtubule arrays respond to cues including light. For example, perception of blue light by hypocotyl epidermal cells leads to reorganization of the microtubule array from a net lateral orientation to a net longitudinal orientation as part of the photomorphogenesis pathway.

Plant cortical microtubule array reorganization requires a set of microtubule regulatory proteins. Along the initial lateral microtubule array, γ-tubulin complexes nucleate new microtubules oriented at an angle from the parental microtubule. Additional γ-tubulin complexes bind these nascent microtubules, leading to the nucleation and polymerization of a set of microtubules arranged orthogonal to the parental lateral array. The orthogonal positioning of microtubules yields microtubule intersections termed crossover sites. The microtubule severing enzyme, katanin, is recruited to nucleation and crossover sites, where the nascent/longitudinally-oriented microtubule is severed, and its minus end stabilized by the protein SPIRAL2 (SPR2) (7–11). The preferential severing and minus-end stabilization of longitudinal microtubules leads to their polymerization and amplification over the parental lateral array. How plant cytoskeletal regulators recognize microtubule minus ends and crossover sites and differentiate lateral versus longitudinal microtubules is poorly understood.

SPR2 (also known as TORTIFOLIA1 and CONVOLUTA) was identified as a factor involved in anisotropic growth in *Arabidopsis*, with mutations leading to right-handed spiral growth (12). Initial investigations demonstrated that SPR2 colocalizes with cortical microtubules, has in vitro microtubule binding activity, affects microtubule dynamics and microtubule array reorientation, and modulates microtubule severing (13–16). Subsequent investigations found that SPR2 family members bind and stabilize the microtubule minus end, both in vivo, and when examined using in vitro microtubule dynamics reconstitution assays (9–11). In metazoans, CAMSAP protein family members bind and regulate microtubule minus ends using a CKK domain (17–21). Higher plants lack CAMSAP proteins, but have members of the plant-specific SPR2 family (22). The domain architecture of SPR2 family members is distinct from CAMSAP proteins, as they contain a predicted N-terminal TOG domain (23–26), a central coiled-coil, and a helical C-terminal domain of unknown structure (Fig 1A). The structure and mechanism of SPR2 microtubule minus end recognition and regulation is a central question in plant cytoskeletal research which requires structural and functional analysis of each conserved domain.

**Fig 1.**
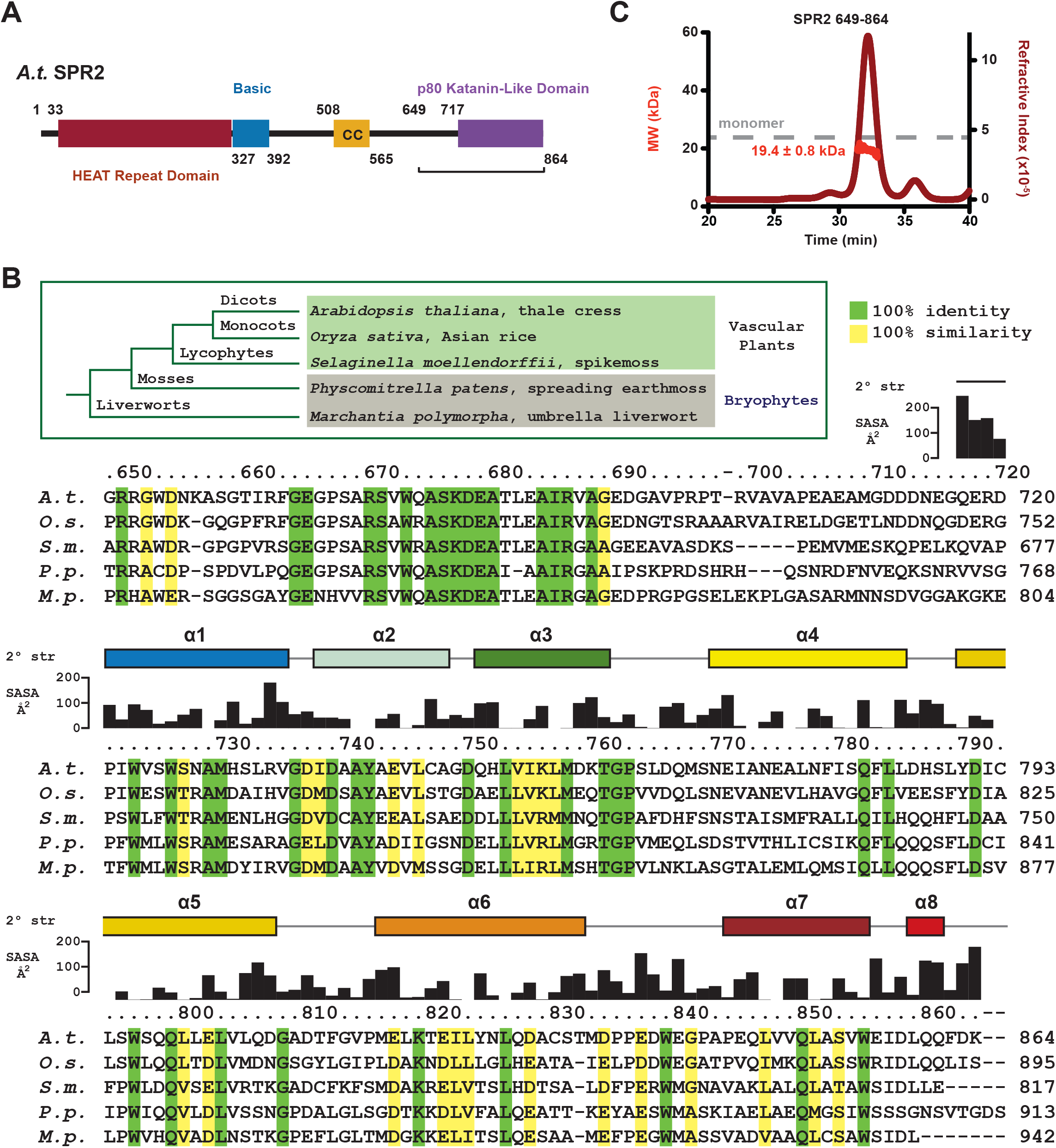
SPR2 contains a conserved C-terminal domain. (A) Domain architecture of SPR2, consisting of an N-terminal HEAT repeat domain, a basic region, a central coiled coil, and a conserved C-terminal domain that structurally resembles the p80 katanin domain involved in p60-p80 katanin heterodimerization. The construct used for crystallization (residues 649-864) is indicated. (B) Sequence alignment of SPR2 homologs from diverse land plants. Conservation is mapped on the sequence alignment as follows: green, 100% identity across species aligned; yellow, 100% similarity across species aligned using the following similarity rubric (LIVM, TSC, RK, NQ, DE, FYW, AG, H, P). Residue numbers are indicated above the alignment for *Arabidopsis thaliana* SPR2, as are secondary structure and residue solvent accessibility, both determined based on the crystal structure of the *A*.*t*. SPR2 C-terminal domain presented here. Inset depicts evolutionary tree of the land plants aligned, including their Latin and common names. (C) SECMALS analysis of SPR2 649-864 (FW 24.2 kDa). Plot shows the elution profile from the size exclusion column as measured using the refractive index (y-axis at right) over time. The experimentally determined mass is plotted in kDa (MW, y-axis at left) over time across the elution peak. The average mass (± standard deviation) is indicated. The dashed gray line indicates the monomeric formula weight.

Here, we explore the structure of the SPR2 C-terminal domain using x-ray crystallography. The aim of the study was to determine the oligomeric state and structure of the SPR2 C-terminal conserved region, compare and contrast the structure with other microtubule associated protein domains, and map conservation on the domain to identify tentative protein-protein interaction sites. We find that the *Arabidopsis thaliana* (*A*.*t*.) SPR2 C-terminal domain is monomeric in solution, and we present the 1.8 Å resolution crystal structure of the domain, which reveals an α-solenoid fold consisting of seven conserved α-helices. Comparison of the SPR2 C-terminal domain structure with similar domain folds from Ge-1 and katanin p80 highlights distinct topological features of SPR2 indicative of distinct function. We identify a conserved face of the SPR2 C-terminal domain likely involved in binding protein partners.

## Materials and methods

### Protein expression and purification

*A*.*t*. SPR2 DNA encoding residues 649-864 was generated using the polymerase chain reaction method (primers: 5’-GGCAGGACCCATATGGGCAGGAGAGGGTGGGATAATAAAGC-3’ and 5’-GCCGAGCCTGAATTCTTACTTGTCGAACTGTTGGAGATCGATTTC-3’) and individually sub-cloned into pET28 (Millipore Sigma, Burlington, MA) using engineered NdeI and EcoRI restriction endonuclease sites, digested, and ligated (New England Biolabs, Ipswich, MA). The construct was transformed into B834(DE3) *E. coli* methionine auxotrophic cells, grown to an optical density at 600 nm of 1.0 in 6l SelenoMet Medium (Molecular Dimensions Limited, Rotherham, UK) containing 50 μg/l kanamycin, 100 μM iron sulfate, and 60 mg/l DL-selenomethionine (Millipore Sigma), the temperature lowered to 20° C, and protein expression induced with 100 μM Isopropyl β-D-1-thiogalactopyranoside for 12 hours. Cells were harvested by centrifugation, and resuspended in 150 ml buffer A (25 mM Tris pH 8.0, 300 mM NaCl, 10 mM imidazole, 0.1% (v/v) β-mercaptoethanol, 5 mM L-methionine) at 4° C, supplemented with DNase (5 μg/ml final concentration, Worthington Biochemical Corp., Lakewood, NJ), lysozyme (10 μg/ml final concentration, Thermo Fisher Scientific, Waltham, MA), and 0.5 mM phenylmethylsulfonyl fluoride (PMSF). Lysis was aided by sonication during which the PMSF final concentration was increased to 1 mM. Lysate was cleared by centrifugation at 23,000 x g for 45 minutes at 4° C. The supernatant was loaded onto a Ni^2+^-NTA column (QIAGEN, Hilden, Germany) and washed with 750 mls of buffer A. Protein was batch eluted with buffer B (buffer A supplemented with 290 mM Imidazole). CaCl_2_ was added to 1 mM final concentration, and 0.1 mg bovine α-thrombin (Haematologic Technologies, Essex Junction, VT) added to proteolytically cleave off the N-terminal His_6_ tag, leaving an N-terminal Gly-Ser-His-Met N-terminal cloning artifact. Protein was dialyzed into buffer A for 24 hrs using 3k MWCO dialysis tubing (Thermo Fisher). Protein was then filtered over a benzamidine-Sepharose column (Cytiva, Marlborough, MA) to remove thrombin. A subsequent Ni^2+^-NTA column was used to remove uncleaved His_6_-tagged protein. Cleaved protein was buffer exchanged into storage buffer (25 mM Tris pH 8.5, 500 mM NaCl, and 0.1% β-mercaptoethanol, 5 mM L-methionine), concentrated using 3 kDa Amicon Ultra Spin Concentrators (MilliporeSigma) to 2.8 mM (68 mg/ml), flash frozen in liquid nitrogen, and stored at -80° C.

### Size exclusion chromatography and multi-angle light scattering

The SPR2 649-864 construct (100 μl of 220 μM protein) was injected onto a Superdex 200 10/300 GL size exclusion column (Cytiva) pre-equilibrated and run in 25 mM Tris pH 8.5, 500 mM NaCl, 0.1% β-mercaptoethanol, 0.2 g/L sodium azide. The protein sample was then directly passed through a Wyatt DAWN HELEOS II light scattering instrument and a Wyatt Optilab rEX refractometer. The light scattering values and the refractive index values were used to calculate the weight-averaged molar mass (M_W_) across the elution peak using the Wyatt Astra V software program (Wyatt Technology Corp., Santa Barbara, CA). Data plots were generated using Prism (GraphPad Software, San Diego, CA). Data shown are representative of duplicate runs.

### Protein crystallization

Selenomethionine (SeMet)-substituted SPR2 (residues 649-864) was crystallized using the hanging drop procedure at 20° C. 2 μl of SPR2 protein at 7 mg/ml was mixed with 2 μl of well solution (1.05 M Ammonium sulfate, 100 mM sodium acetate pH 4.6), placed on a silanized glass coverslip, and used to seal a chamber containing 1 ml of the well solution. Crystals formed overnight and continued to grow over the course of a week. Single crystals were harvested, transferred to FOMBLIN Y (MilliporeSigma), flash frozen in liquid nitrogen, and stored in liquid nitrogen.

### Data collection, structure determination, refinement, and analysis

A selenium SAD peak data set at 12,661.01 eV (λ = 0.9792603 Å) was collected on a single crystal to a resolution of 1.8 Å. Diffraction data were collected at the Advanced Photon Source beamline 22-ID at 100 K in 0.5° oscillations, across 360°. Crystals belong to the P2_1_2_1_2_1_ space group with one molecule in the asymmetric unit. Data were indexed, integrated, and scaled using HKL2000(27). Selenium sites were identified and used to generate initial density-modified electron density maps using PHENIX AutoSol(28). Initial models were built using AutoBuild (PHENIX), followed by reiterative manual building in Coot and refinement using phenix.refine(28,29). The SeMet-substituted structure was refined against an MLHL target function. The free *R* used 10% of the data randomly excluded from refinement. Information regarding data statistics, model building, and refinement is presented in Table 1. Electrostatics was calculated using APBS(30). Protein Data Bank (PDB) structure similarity searches were performed using the Dali server(31). Pairwise structural alignments and rmsd values were calculated using the PDBeFold server(32). Solvent accessibility was calculated using the PDBePISA server(33). Structure figures were generated using PyMOL (Schrödinger, New York, NY). Sequence alignments were generated using Clustal Omega(34) followed by manual adjustments. Secondary structure prediction used the Jpred4 server(35).

**Table 1.** Crystallographic data processing and refinement statistics.

|  |  |
| --- | --- |
| **Crystal** | **A.t. SPR2 residues 649-864** |
| **Data Collection** |  |
| Wavelength (Å) | 0.9792603 |
| Space group | P 2<sub>1</sub> 2<sub>1</sub> 2<sub>1</sub> |
| Cell dimensions: a, b, c (Å) | 35.5, 47.7, 111.1 |
| Resolution (Å) | 50.00-1.80 (1.86-1.80) |
| # Reflections: Measured / Unique | 187,724 (10,242) / 18,011 (1679) |
| Completeness (%) | 98.3 (92.9) |
| Mean redundancy | 10.4 (6.1) |
| <I/σI> | 21.8 (6.4) |
| R<sub>sym</sub> | 0.098 (0.263) |
| CC1/2 | 0.994 (0.973) |
| CC* | 0.998 (0.993) |
| **Refinement** |  |
| Resolution (Å) | 33.80-1.80 (1.85-1.80) |
| R/ R<sub>free</sub> (%) | 18.6 (20.9) / 21.7 (28.2) |
| # Reflections, R/R<sub>free</sub> | 16121 (1113) / 1792 (123) |
| Total atoms: Protein / Water | 1173 / 95 |
| Residues modeled | 717-864 |
| Stereochemical ideality (rmsd): Bonds (Å) / Angles (°) | 0.011 / 1.323 |
| Ramachandran Analysis: Favored / Allowed (%) | 99.3 / 0.7 |
| PDB Accession Code | 8F8N |

Values in parentheses indicate statistics for the highest-resolution shell.

## Results and discussion

### The SPR2 C-terminal region is highly conserved across land plants

To gain insight into the structure of the SPR2 C-terminal region, we aligned SPR2 homologs from diverse land plants including bryophytes such as liverwort and spreading earthmoss, and vascular plants such as spikemoss, Asian rice, and thale cress (*A*.*t*.) (Fig 1B). SPR2 homologs aligned well over this C-terminal region, with a cluster of sequence identity corresponding to *A*.*t*. SPR2 residues 664-689, and across the region spanning 723-855. A segment of low identity and variable length bridges these two regions across the species aligned. Overall, across the ≥ 450 million years of divergence represented by these species (36,37), their SPR2 homologs have about 19% sequence identity across the C-terminal region. Based on this conservation, we cloned a SPR2 construct embodying residues 649-864, expressed the construct in *E. coli*, and purified the protein to homogeneity.

To determine whether the SPR2 C-terminal region is monomeric or oligomeric, we analyzed the construct using size exclusion chromatography multi angle light scattering (SECMALS) (Fig 1C). SPR2 649-864 eluted as one main peak with an experimentally determined mass of 19.4 ± 0.8 kDa, similar to the monomeric formula weight of 24.2 kDa, indicating that the construct is monomeric at the concentration examined.

### The SPR2 C-terminal domain is a conserved, 7-helix α-solenoid

To gain insight into the architecture of the SPR2 C-terminal region, we crystallized the *A*.*t*. SPR2 649-864 construct and determined its three-dimensional structure. We expressed, purified, and crystallized SeMet-substituted SPR2 649-864 and collected a single wavelength anomalous diffraction (SAD) data set at the selenium peak to 1.8 Å resolution. The crystal belonged to the space group P2_1_2_1_2_1_, with one SPR2 649-864 molecule in the asymmetric unit, and a solvent content of 35% (Fig 2). Selenium sites were identified and used to phase the structure, yielding clear, interpretable electron density (Final 2mF_o_-DF_c_ electron density shown in Fig 2B,D), for which residues 717-849 were modeled. No electron density was apparent for the highly conserved region spanning residues 664-689. The final model was refined to an R value of 18.6%, and a R_free_ value of 21.7%. See Table 1 for crystallographic and refinement statistics.

**Fig 2.**
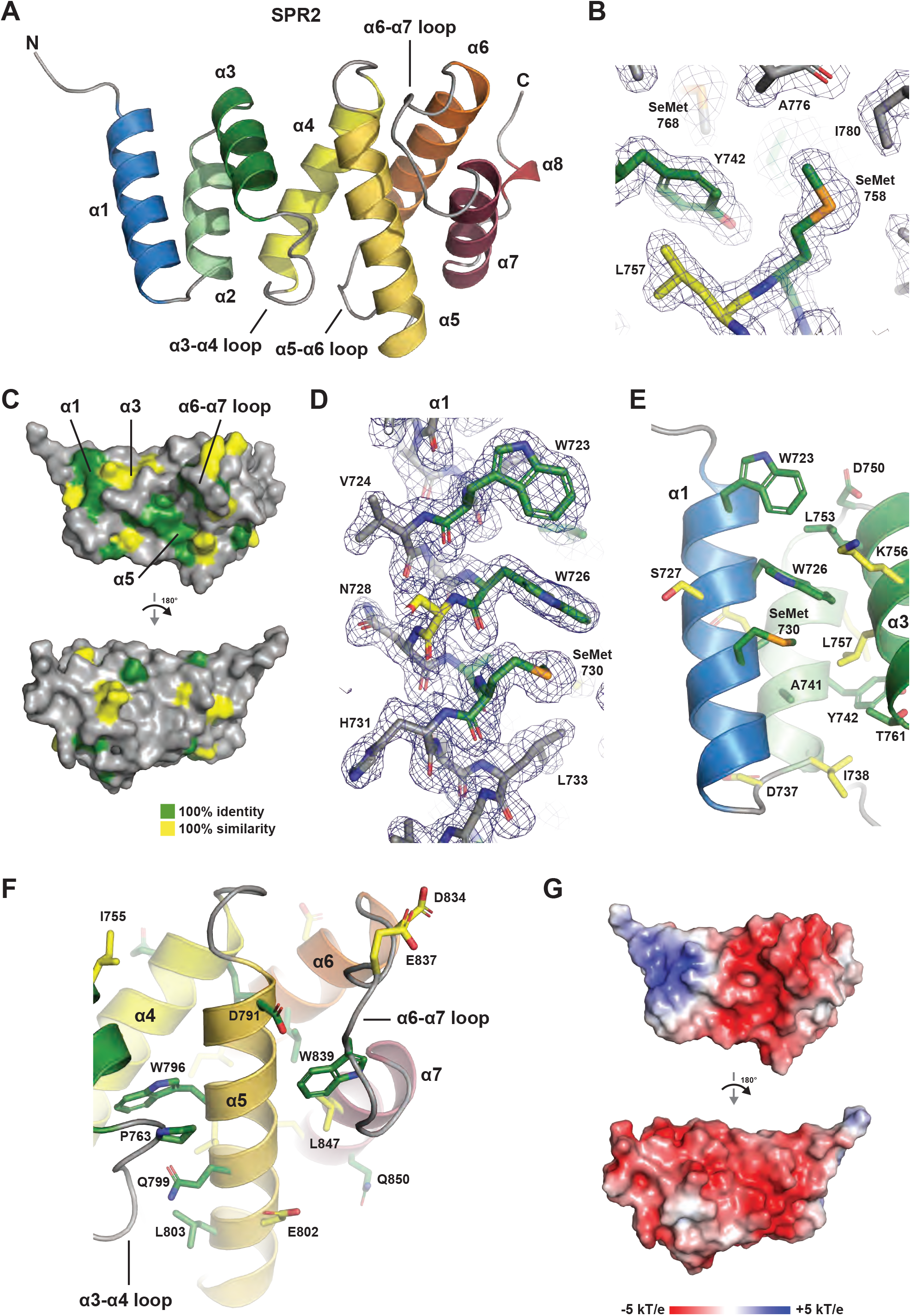
The SPR2 C-terminal domain is an α-solenoid helix-turn-helix domain containing seven helices. (A) Structure of the *A*.*t*. SPR2 C-terminal domain shown in cartoon format. The seven core helices of the domain (α1-α7) are colored across the spectrum. A final helix, α8, packs against the domain, but is not conserved across SPR2 homologs, and thus is not considered part of the core domain. (B) Conserved residues in the α2-α3 region contribute the domain’s hydrophobic core. Residues are shown in stick format, with conservation colored as in Fig 1B, with final 2mF_o_-DF_c_ electron density shown in blue, contoured at 1.0 σ. Two SeMet residues used in phasing are indicated. (C) The SPR2 C-terminal domain shown in surface representation, with conservation from Fig 1B mapped on the surface (green: 100% identity; yellow: 100% similarity). Orientation at top as shown in A, orientation below after a 180° rotation about the y-axis. (D) Conserved, hydrophobic, surface exposed determinants of the α1 helix (W723, W726, and SeMet730) are shown in stick format. Conservation is colored as in Fig 1B, with final 2mF_o_-DF_c_ electron density shown in blue, contoured at 1.0 σ. (E) View of conserved residues in the α1 – α3 region, highlighting the surface exposed hydrophobic and basic nature of region. Backbone shown in cartoon format, colored as in A, with conserved residues shown in stick format, colored as in Fig 1B. (F) View of conserved residues in the α3 – α7 region, highlighting the conserved, surface exposed residues along α5, residue P763 of the α3-α4 loop, as well as residue W839 of the α6-α7 loop, which is positioned in a pocket on the side of the domain. Backbone shown in cartoon format, colored as in A, with conserved residues shown in stick format, colored as in Fig 1B. (G) Electrostatic surface potential mapped on the SPR2 C-terminal domain structure, views oriented as in C.

SPR2 residues 717-849 form a right-handed α-solenoid helix-turn-helix structure, composed of seven conserved α-helices (α1-α7) (Fig 2A). The dimensions of the domain are approximately 45 Å along the axis of the solenoid, 35 Å high, and 30 Å wide. A short helix, α8, packs against the domain, but this segment is not conserved across SPR2 homologs (Fig 1B) and is thus not considered part of the domain’s core structure. Six of the seven α-helices form anti-parallel helix-turn-helix pairs that pack against one another. A number of helix-turn-helix motifs form α-solenoid structures including <u>H</u>untingtin, <u>E</u>longation factor 3, protein phosphatase 2<u>A</u>, <u>T</u>OR1 (HEAT), armadillo (ARM), and FANC repeats (38). Of these repeats, the SPR2 C-terminal domain helix-turn-helix motifs are structurally most similar to FANC repeats, as the helices are relatively straight, and lack a canonical kink present in the first helix of a HEAT repeat, or the additional N-terminal helix of an ARM repeat. The kink in the HEAT repeat structure is due to a proline residue in the first helix, while the separate N-terminal helix of ARM repeats is delineated by a position specific glycine and proline residue that position the N-terminal helix orthogonal to the axes of the subsequent two helices. The helix-turn-helix motifs of the SPR2 C-terminal domain lack these specific proline and glycine residues. The two helices in each pair form a hydrophobic interface between each other, and with the flanking helices, collectively forming a hydrophobic core that runs along the axis of the α-solenoid (Fig 2B). The loops between helices vary in length, both within a helix-turn-helix motif, and between these motifs. Extended ordered loops of conserved length include the α3-α4 loop, the α5-α6 loop, and the α6-α7 loop (Fig 2A).

Conservation, as contoured in Fig 1B, maps primarily to one face of the domain, with a high degree of identity conserved over ≥ 450 million years of evolution (Fig 2C). Key contributions to this conserved face come from surface-exposed hydrophobic residues on α1 (Fig 2D,E), including W723, W726, M730, and a cluster of hydrophobic residues on the α3-α4 loop - α5 interface, including P763, which stacks against W796, as well as L803 (Fig 2F). The α6-α7 loop forms an extensive projection from the domain that packs against α5, forming a hydrophobic pocket involving W839 (Fig 2F). The domain has a net negative charge (Fig 2G). On the conserved face of the domain, charge is partitioned, with a basic patch localized to the α1-α3 region (Fig 2G). Collectively, conservation mapping suggests that the domain face formed by α1, α3, α5, and the α6-α7 loop is likely to constitute a functional surface, potentially for protein-protein interactions, mediated by both hydrophobic and electrostatic interactions.

### The SPR2 C-terminal α-solenoid domain is structurally homologous to the C-terminal domains of Ge-1 and the katanin p80 subunit

To determine whether the SPR2 C-terminal domain (residues 717-849) is structurally homologous to other protein structures, we used the Dali server (31) to search the PDB, which identified two highly homologous domain structures: the C-terminal domains from Ge-1 and the katanin p80 subunit. Ge-1 is part of the mRNA 5’ decapping complex, and is involved in localizing the complex to the P-body (39,40). The *Drosophila melanogaster* Ge-1 structure (PDB accession code 2VXG, chain B (41)) structurally aligns well with the SPR2 C-terminal domain (Z-score 10.7, 2.6 Å rmsd over 119 Cα atoms, 16% sequence identity) (Fig 3A-C). The Ge-1 C-terminal domain consists of a core eight α-helices. Ge-1 helices α1-α3 and α5-α8 correspond to SPR2 helices α1-α7 respectively, Ge-1 has a unique α4 helix, positioned perpendicular to α3, that bridges the second and third paired helices, and is followed by a disordered loop. Additional key structural differences between SPR2 and Ge-1 include (using SPR2 nomenclature) the SPR2 α4-α5 loop, the SPR2 α5-α6 loop (for which the corresponding loop in Ge-1 is flanked by a shorter N-terminal helix, and a longer, and kinked C-terminal helix), and the SPR2 α6-α7 loop (which is extended in SPR2, and in Ge-1 is flanked by a longer C-terminal helix) (Fig 3C). In contrast to SPR2, Ge-1 conservation maps primarily to the opposite face of the domain, including residues on Ge-1 α5 (structurally equivalent to SPR2 α4), and a conserved arginine on Ge-1 α8, which when mutated (R1340E), affects the ability of Ge-1 to localize to P-bodies (41). The Ge-1 C-terminal domain also has a distinct, net basic electrostatic surface potential (Fig 3D). Overall, the Ge-1 C-terminal domain, while similar to SPR2 in fold, has distinct structural attributes, surface conservation and electrostatics, suggesting that the common fold is involved in distinct, non-overlapping functions for these proteins.

**Fig 3.**
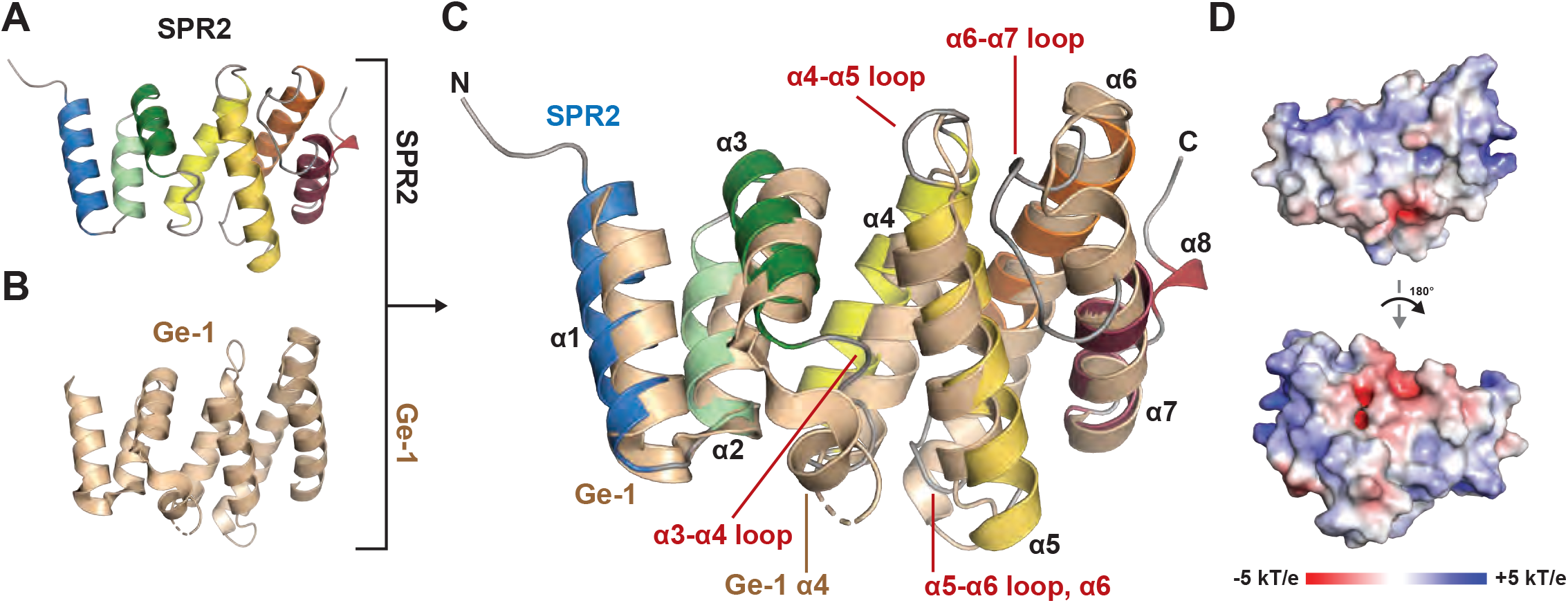
The SPR2 C-terminal domain is structurally similar to the C-terminal domain from the mRNA 5’-decapping factor, Ge-1. (A) Structure of the *A*.*t*. SPR2 C-terminal domain, colored as in Fig 2A, shown in cartoon format. (B) Structure of the *Drosophila melanogaster* Ge-1 C-terminal domain (colored wheat, shown in cartoon format (PDB accession code: 2VXG, Chain A (41)). (C) Structural alignment of the SPR2 C-terminal domain and the *D*.*m*. Ge-1 C-terminal domain from 2VXG(41), oriented as in A and B. Major differences in domain architecture are labeled in red. Labels denote SPR2 secondary structure elements unless otherwise noted. (D) Electrostatic surface potential mapped on the Ge-1 C-terminal domain structure 2VXG (Chain A(41)), top view oriented as in A, bottom view after a 180° rotation about the y-axis.

The second hit from the Dali server (31) search of the PDB we performed was the regulatory p80 subunit of the katanin microtubule severing enzyme. Katanin consists of a catalytic p60 subunit and a non-catalytic, regulatory p80 subunit (42). The p60 subunit has an AAA+ domain that hexamerizes into a lock washer structure that pulls, in an ATP-hydrolysis-dependent manner, on a microtubule lattice β-tubulin tail. Katanin extracts the tubulin subunit from the lattice, leading either to repair (incorporation of GTP-bound tubulin), or lattice destabilization and severing (43–47). N-terminal to AAA+ domain is a microtubule-interacting and -trafficking (MIT) domain, which heterodimerizes with the Katanin p80 C-terminal domain (42,48–50). As SPR2 and katanin are both involved in reorientation of the plant microtubule array, we compare and contrast the SPR2 and p80 C-terminal domain structures in detail.

The SPR2 C-terminal domain aligns well with the p80 C-terminal domain structure, which was determined in complex with the p60 katanin MIT domain (Z-score 9.8, 3.6 Å rmsd over 120 Cα atoms, 13% sequence identity, compared with PDB accession code 5NBT, chain C (50)) (Fig 4A-C). The p80 C-terminal domain consists of the seven core α-helices that align well with the SPR2 C-terminal domain α-helices. However, we do note the following structural differences. First, p80 katanin α1 has a long N-terminal extension that is involved in binding the p60 MIT domain. Second, the p80 katanin α3-α4 region diverges as follows: p80 α3 helix is extended relative to SPR2 α3, and the p80 α4 N-terminal region is kinked due to a proline residue in the middle of α4 that contrasts with SPR2’s straight α4 helix. Collectively, these differences position the p80 α3-α4 loop in a conformation distinct from the SPR2 α3-α4 loop. Third, p80 katanin α6 is shifted relative to SPR2 α6 (along the helical axis), and the loops that flank p80 α6 are disordered. While the SPR2 α5-α6 loop is ordered, the p80 α5-α6 loop is much longer and includes 15 residues not ordered in the structure. Similarly, the SPR2 α6-α7 loop forms an ordered 11-residue structure that packs against α5, while the 11-residue p80 α6-α7 loop could not be modeled over 10 of the 11 residues.

**Fig 4.**
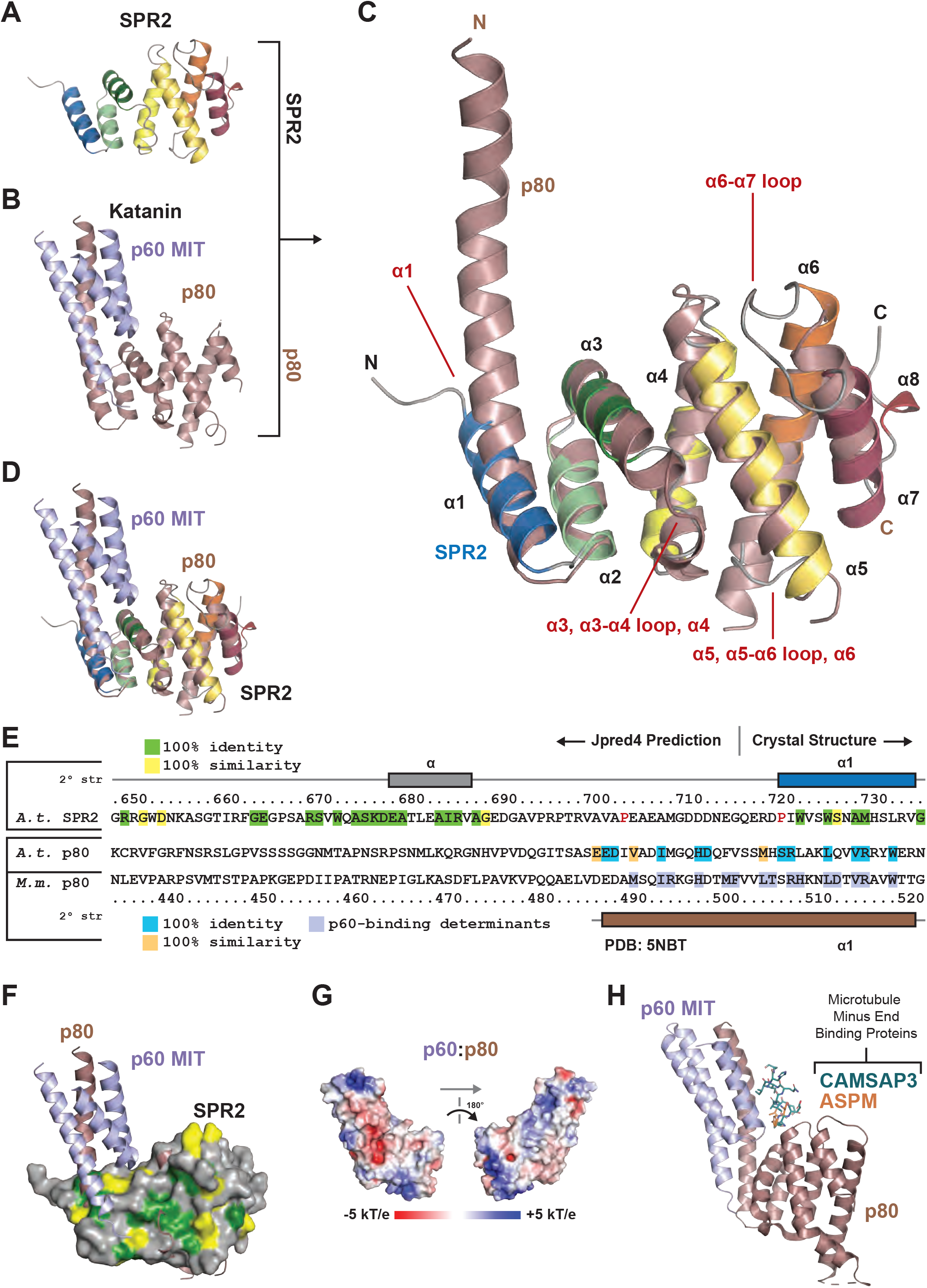
The SPR2 C-terminal domain is structurally similar to the katanin p80 C-terminal domain that heterodimerizes with the katanin p60 MIT domain. (A) Structure of the *A*.*t*. SPR2 C-terminal domain, colored as in Fig 2A, shown in cartoon format. (B) Structure of the mouse katanin p60:p80 heterodimeric complex, involving the p60 MIT domain (colored light blue) and the p80 C-terminal domain (colored rose gold)(PDB accession code: 5NBT(50)). (C) Structural alignment of the SPR2 C-terminal domain and the katanin p80 C-terminal domain from 5NBT(50), oriented as in A and B. Major differences in domain architecture are labeled in red. Labels denote SPR2 secondary structure elements. (D) Structural alignment of the SPR2 C-terminal domain and the Katanin p60:p80 heterodimer from 5NBT(50), oriented as in A and B. (E) Sequence alignment of the N-terminal region, including α1, from the *A*.*t*. SPR2 C-terminal domain and the *A*.*t*. and Mus musculus (*M*.*m*.) p80 katanin C-terminal domain. Conservation across SPR2 homologs from Fig 1B is indicated on the SPR2 sequence. *M*.*m*. p80 residues involved in contacts with p60 are highlighted in light blue. Residues conserved between *M*.*m*. p80 and *A*.*t*. p80 over the region modeled in the 5NBT (50) structure are highlighted dark cyan (100% identity) and light orange (100% similarity) on the *A*.*t*. p80 sequence. Residue numbers are for *A*.*t*. SPR2 (above the alignment) and *M*.*m*. p80 (below the alignment). Secondary structure is indicated above for SPR2 based on the crystal structure (residues 717-736), and predicted using Jpred4 (for residues 649-716, which were not ordered in the crystal structure), while the secondary structure for *M*.*m*. p80 is shown below based on the 5NBT structure (50). Proline residues in the SPR2 sequence that are N-terminal to α1 and within the equivalent span that constitutes α1 in the *M*.*m*. p80 structure are colored red. (F) Structural alignment of the SPR2 C-terminal domain and the katanin p60:p80 heterodimer from 5NBT(50), oriented as D, with SPR2 shown in surface representation with conservation mapped as in Fig 2C. (G) Electrostatic surface potential mapped on the katanin p60:p80 heterodimer structure (50) (left image oriented as in B, right image after a 180° rotation about the y-axis). (H) Structural alignment of the mouse katanin p60:p80 heterodimerization module in complex with the microtubule minus end-binding proteins: Abnormal spindle-like microcephaly-associated protein homolog (ASPM, PDB accession code 5LB7 (49), shown in stick format, colored orange) and CAMSAP3 (PDB accession code 5OW5 (51), shown in stick format, colored dark teal). The p60 and p80 chains are only shown from the 5LB7 structure for simplicity.

The katanin p60 and p80 subunits form an extensive interaction along the length of p80 α1 (50). As the interaction with p60 likely stabilized the extended α1 helix, we investigated whether the sequence N-terminal to the SPR2 α1 helix modeled in our structure, might contain homology to the p80 subunit’s α1 p60-binding determinants, and whether this might suggest an ability of the SPR2 C-terminal domain to directly bind p60. Using the structural alignment as shown in Fig 4C, inclusion of the p60 MIT domain from the 5NBT structure (50) leads to steric clash between the MIT domain and residues on SPR2 α1 and α3 (Fig 4D). While katanin p60:p80 interactions are primarily hydrophobic, two key hydrophobic residues in p80 α3 correspond with lysine residues in the SPR2 structure, which we anticipate would prohibit p60 and SPR2 from engaging in a similar mode as observed in the katanin p60:p80 heterodimer structure (50). While many p80 α1 residues involved in p60 binding are conserved between mouse p80 and *A*.*t*. p80, few of these residues are found in SPR2 (Fig 4E). Of note, secondary structure prediction using Jpred4 (35) predicts a disordered region over the SPR2 span equivalent to the p80 α1 N-terminal extension (Fig 4E). This span of SPR2 also includes two proline residues, which would be predicted to compromise formation of a straight helix over the span (Fig 4E). While there is a conserved region 32 residues N-terminal to SPR2 α1, this region has no similarity to p80. Katanin p60 does engage the katanin p80 C-terminal domain over a region that corresponds to a conserved site on the SPR2 C-terminal domain structure involving residues from α1 and α3 (Fig 4F). This suggests that similar regions of the SPR2 and p80 C-terminal domains may be involved in protein-protein interactions. The katanin p60:p80 complex has significant basic electrostatic patches (Fig 4G), that align with the complex’s ability to bind the negatively-charged microtubule exterior (48–50). This contrasts with the highly acidic electrostatics of the SPR2 C-terminal domain (Fig 2G). Overall, while the SPR2 and katanin p80 C-terminal domains are structurally similar, they have distinct architectural differences, conservation, and electrostatics. Based on these differences, we do not anticipate that SPR2 engages katanin p60 using a p80-binding mode.

## Conclusion

We experimentally determined the structure of the SPR2 conserved C-terminal domain, revealing a domain fold found in the mRNA de-capping component, Ge-1, and the katanin microtubule severing enzyme regulatory p80 subunit. The SPR2 structure has distinct conformations, conservation, and electrostatics that set it apart from Ge-1 and p80 katanin, suggesting that its function is also distinct. Interestingly, both SPR2 and katanin play central roles in the reorganization of the microtubule array in plants in response to blue light. Katanin is recruited to microtubule crossover sites, where it severs microtubules oriented in the longitudinal array, thereby amplifying the number of microtubules in the longitudinal array (7). SPR2 recognizes and stabilizes microtubule minus ends, which is critical to prevent depolymerization of the longitudinal array (9–11). How katanin is recruited to microtubule crossover sites and specifically cleaves the longitudinally-oriented microtubule remains to be fully determined (8), as is the mechanism by which SPR2 specifically binds and stabilizes the microtubule minus end. Our structural work reveals an interesting evolutionary relation between SPR2 and katanin p80, in that they have a common structural domain. While we do not anticipate binding between SPR2 and katanin p60 in a mode analogous to the katanin p60:p80 complex (49,50), whether SPR2 and katanin interact remains to be experimentally determined. Interestingly, the mammalian katanin p60:p80 complex uses a common site to bind CAMSAP3 (51) and ASPM (49), two proteins that directly recognize and bind the microtubule minus end (Fig 4H), highlighting the potential evolutionary functional convergence of the katanin p80/SPR2 domain as a determinant at the nexus of microtubule severing and microtubule minus end localization. The SPR2 C-terminal domain structure lays a foundation upon which its role in the regulation of microtubule minus end dynamics and array reorientation can be investigated.

## Acknowledgments

We thank the staff at the Argonne National Laboratory Advanced Photon Source SER-CAT beamline 22-ID for support. We thank Dr. Ashutosh Tripathy for assistance with SECMALS and acknowledge use of the UNC Macromolecular Interactions Facility.

## References

1. Mitchison T, Kirschner M. Dynamic instability of microtubule growth. Nature. 1984 Nov 15;312(5991):237–42.

2. Desai A, Mitchison TJ. Microtubule polymerization dynamics. Annu Rev Cell Dev Biol. 1997;13:83– 117.

3. Baskin TI. On the alignment of cellulose microfibrils by cortical microtubules: a review and a model. Protoplasma. 2001;215(1–4):150–71.

4. Ehrhardt DW, Shaw SL. Microtubule dynamics and organization in the plant cortical array. Annu Rev Plant Biol. 2006;57:859–75.

5. Wasteneys GO. Microtubule organization in the green kingdom: chaos or self-order? J Cell Sci. 2002 Apr 1;115(Pt 7):1345–54.

6. Paredez AR, Somerville CR, Ehrhardt DW. Visualization of cellulose synthase demonstrates functional association with microtubules. Science. 2006 Jun 9;312(5779):1491–5.

7. Lindeboom JJ, Nakamura M, Hibbel A, Shundyak K, Gutierrez R, Ketelaar T, et al. A Mechanism for Reorientation of Cortical Microtubule Arrays Driven by Microtubule Severing. Science. 2013 Dec 6;342(6163):1245533.

8. Yagi N, Kato T, Matsunaga S, Ehrhardt DW, Nakamura M, Hashimoto T. An anchoring complex recruits katanin for microtubule severing at the plant cortical nucleation sites. Nat Commun. 2021 Jun 17;12(1):3687.

9. Nakamura M, Lindeboom JJ, Saltini M, Mulder BM, Ehrhardt DW. SPR2 protects minus ends to promote severing and reorientation of plant cortical microtubule arrays. J Cell Biol. 2018 Mar 5;217(3):915–27.

10. Leong SY, Yamada M, Yanagisawa N, Goshima G. SPIRAL2 Stabilises Endoplasmic Microtubule Minus Ends in the Moss Physcomitrella patens. Cell Struct Funct. 2018 Mar 28;43(1):53–60.

11. Fan Y, Burkart GM, Dixit R. The Arabidopsis SPIRAL2 Protein Targets and Stabilizes Microtubule Minus Ends. Curr Biol. 2018 Mar 19;28(6):987-994.e3.

12. Furutani I, Watanabe Y, Prieto R, Masukawa M, Suzuki K, Naoi K, et al. The SPIRAL genes are required for directional control of cell elongation in Aarabidopsis thaliana. Development. 2000 Oct;127(20):4443–53.

13. Buschmann H, Fabri CO, Hauptmann M, Hutzler P, Laux T, Lloyd CW, et al. Helical growth of the Arabidopsis mutant tortifolia1 reveals a plant-specific microtubule-associated protein. Curr Biol. 2004 Aug 24;14(16):1515–21.

14. Shoji T, Narita NN, Hayashi K, Hayashi K, Asada J, Hamada T, et al. Plant-specific microtubule-associated protein SPIRAL2 is required for anisotropic growth in Arabidopsis. Plant Physiol. 2004 Dec;136(4):3933–44.

15. Wightman R, Chomicki G, Kumar M, Carr P, Turner SR. SPIRAL2 determines plant microtubule organization by modulating microtubule severing. Curr Biol. 2013 Oct 7;23(19):1902–7.

16. Yao M, Wakamatsu Y, Itoh TJ, Shoji T, Hashimoto T. Arabidopsis SPIRAL2 promotes uninterrupted microtubule growth by suppressing the pause state of microtubule dynamics. J Cell Sci. 2008 Jul 15;121(Pt 14):2372–81.

17. Meng W, Mushika Y, Ichii T, Takeichi M. Anchorage of Microtubule Minus Ends to Adherens Junctions Regulates Epithelial Cell-Cell Contacts. Cell. 2008 Nov 28;135(5):948–59.

18. Goodwin SS, Vale RD. Patronin regulates the microtubule network by protecting microtubule minus ends. Cell. 2010 Oct 15;143(2):263–74.

19. Nagae S, Meng W, Takeichi M. Non-centrosomal microtubules regulate F-actin organization through the suppression of GEF-H1 activity. Genes Cells. 2013 May;18(5):387–96.

20. Tanaka N, Meng W, Nagae S, Takeichi M. Nezha/CAMSAP3 and CAMSAP2 cooperate in epithelial-specific organization of noncentrosomal microtubules. Proc Natl Acad Sci U S A. 2012 Dec 4;109(49):20029–34.

21. Atherton J, Luo Y, Xiang S, Yang C, Rai A, Jiang K, et al. Structural determinants of microtubule minus end preference in CAMSAP CKK domains. Nat Commun. 2019 Nov 20;10(1):5236.

22. Akhmanova A, Steinmetz MO. Microtubule minus-end regulation at a glance. Journal of Cell Science. 2019 Jun 7;132(11):jcs227850.

23. Hamada T. Lessons from in vitro reconstitution analyses of plant microtubule-associated proteins. Front Plant Sci. 2014;5:409.

24. Haikonen T, Rajamäki ML, Valkonen JPT. Interaction of the microtubule-associated host protein HIP2 with viral helper component proteinase is important in infection with potato virus A. Mol Plant Microbe Interact. 2013 Jul;26(7):734–44.

25. Slep KC, Vale RD. Structural basis of microtubule plus end tracking by XMAP215, CLIP-170, and EB1. Mol Cell. 2007 Sep 21;27(6):976–91.

26. Al-Bassam J, Larsen NA, Hyman AA, Harrison SC. Crystal structure of a TOG domain: conserved features of XMAP215/Dis1-family TOG domains and implications for tubulin binding. Structure. 2007 Mar;15(3):355–62.

27. Otwinowski Z, Minor W. Processing of X-ray diffraction data collected in oscillation mode. Methods Enzymol. 1997;276:307–26.

28. Adams PD, Afonine PV, Bunkóczi G, Chen VB, Davis IW, Echols N, et al. PHENIX: a comprehensive Python-based system for macromolecular structure solution. Acta Crystallogr D Biol Crystallogr. 2010 Feb;66(Pt 2):213–21.

29. Emsley P, Lohkamp B, Scott WG, Cowtan K. Features and development of Coot. Acta Crystallogr D Biol Crystallogr. 2010 Apr;66(Pt 4):486–501.

30. Baker NA, Sept D, Joseph S, Holst MJ, McCammon JA. Electrostatics of nanosystems: Application to microtubules and the ribosome. Proceedings of the National Academy of Sciences. 2001 Aug 28;98(18):10037–41.

31. Holm L. Dali server: structural unification of protein families. Nucleic Acids Res. 2022 May 24;gkac387.

32. Krissinel E, Henrick K. Secondary-structure matching (SSM), a new tool for fast protein structure alignment in three dimensions. Acta Cryst D. 2004 Dec 1;60(12):2256–68.

33. Krissinel E, Henrick K. Inference of macromolecular assemblies from crystalline state. J Mol Biol. 2007 Sep 21;372(3):774–97.

34. Madeira F, Pearce M, Tivey ARN, Basutkar P, Lee J, Edbali O, et al. Search and sequence analysis tools services from EMBL-EBI in 2022. Nucleic Acids Res. 2022 Apr 1;gkac240.

35. Drozdetskiy A, Cole C, Procter J, Barton GJ. JPred4: a protein secondary structure prediction server. Nucleic Acids Research. 2015 Jul 1;43(W1):W389–94.

36. Bowman JL. Walkabout on the long branches of plant evolution. Curr Opin Plant Biol. 2013 Feb;16(1):70–7.

37. Bowman JL, Kohchi T, Yamato KT, Jenkins J, Shu S, Ishizaki K, et al. Insights into Land Plant Evolution Garnered from the Marchantia polymorpha Genome. Cell. 2017 Oct 5;171(2):287-304.e15.

38. Nookala RK, Hussain S, Pellegrini L. Insights into Fanconi Anaemia from the structure of human FANCE. Nucleic Acids Research. 2007 Mar 1;35(5):1638–48.

39. Fenger-Grøn M, Fillman C, Norrild B, Lykke-Andersen J. Multiple processing body factors and the ARE binding protein TTP activate mRNA decapping. Mol Cell. 2005 Dec 22;20(6):905–15.

40. Yu JH, Yang WH, Gulick T, Bloch KD, Bloch DB. Ge-1 is a central component of the mammalian cytoplasmic mRNA processing body. RNA. 2005 Dec;11(12):1795–802.

41. Jinek M, Eulalio A, Lingel A, Helms S, Conti E, Izaurralde E. The C-terminal region of Ge-1 presents conserved structural features required for P-body localization. RNA. 2008 Oct;14(10):1991–8.

42. Hartman JJ, Mahr J, McNally K, Okawa K, Iwamatsu A, Thomas S, et al. Katanin, a microtubule-severing protein, is a novel AAA ATPase that targets to the centrosome using a WD40-containing subunit. Cell. 1998 Apr 17;93(2):277–87.

43. McNally FJ, Vale RD. Identification of katanin, an ATPase that severs and disassembles stable microtubules. Cell. 1993 Nov 5;75(3):419–29.

44. Hartman JJ, Vale RD. Microtubule disassembly by ATP-dependent oligomerization of the AAA enzyme katanin. Science. 1999 Oct 22;286(5440):782–5.

45. Zehr E, Szyk A, Piszczek G, Szczesna E, Zuo X, Roll-Mecak A. Katanin spiral and ring structures shed light on power stroke for microtubule severing. Nat Struct Mol Biol. 2017 Sep;24(9):717–25.

46. Zehr EA, Szyk A, Szczesna E, Roll-Mecak A. Katanin Grips the β-Tubulin Tail through an Electropositive Double Spiral to Sever Microtubules. Dev Cell. 2020 Jan 6;52(1):118-131.e6.

47. Vemu A, Szczesna E, Zehr EA, Spector JO, Grigorieff N, Deaconescu AM, et al. Severing enzymes amplify microtubule arrays through lattice GTP-tubulin incorporation. Science. 2018 24;361(6404).

48. McNally KP, Bazirgan OA, McNally FJ. Two domains of p80 katanin regulate microtubule severing and spindle pole targeting by p60 katanin. Journal of Cell Science. 2000 May 1;113(9):1623–33.

49. Jiang K, Rezabkova L, Hua S, Liu Q, Capitani G, Altelaar AFM, et al. Microtubule minus-end regulation at spindle poles by an ASPM-katanin complex. Nat Cell Biol. 2017 May;19(5):480–92.

50. Rezabkova L, Jiang K, Capitani G, Prota AE, Akhmanova A, Steinmetz MO, et al. Structural basis of katanin p60:p80 complex formation. Sci Rep. 2017 Nov 2;7(1):14893.

51. Jiang K, Faltova L, Hua S, Capitani G, Prota AE, Landgraf C, et al. Structural Basis of Formation of the Microtubule Minus-End-Regulating CAMSAP-Katanin Complex. Structure. 2018 Mar 6;26(3):375-382.e4.

